## Supplementary material for "Assessing the performance of genome-wide association studies for predicting disease risk": S1 Table

| GWAS-ROCS ID | Condition/Phenotype | Pawitan et al. Heritability |
|---|---|---|
| GR00001 | Abdominal aortic aneurysm | 0.004162347 |
| GR00002 | Abdominal aortic aneurysm | 0.021762399 |
| GR00003 | Acne (severe) | 0.042017796 |
| GR00004 | Acute lymphoblastic leukemia (childhood) | 0.103370717 |
| GR00005 | Acute lymphoblastic leukemia (childhood) | 0.087244098 |
| GR00006 | Acute lymphoblastic leukemia (childhood) | 0.008476861 |
| GR00007 | Acute-on-chronic liver failure in hepatitis B | 0.027527524 |
| GR00008 | Adolescent idiopathic scoliosis | 0.00985719 |
| GR00009 | Adolescent idiopathic scoliosis | 0.06022384 |
| GR00010 | Age-related macular degeneration | 0.284701365 |
| GR00011 | Age-related macular degeneration | 0.058090794 |
| GR00012 | Age-related macular degeneration | 0.272397476 |
| GR00013 | Age-related macular degeneration | 0.021835469 |
| GR00014 | Age-related macular degeneration | 0.196292911 |
| GR00015 | Age-related macular degeneration | 0.064377847 |
| GR00016 | Age-related macular degeneration | 0.071985892 |
| GR00017 | Aggressive periodontitis | 0.042284365 |
| GR00018 | Allergic disease | 0.022049233 |
| GR00019 | Allergic sensitization | 0.029839602 |
| GR00020 | Alzheimer's disease | 0.008889895 |
| GR00021 | Alzheimer's disease | 0.009514793 |
| GR00022 | Alzheimer's disease | 0.004295769 |
| GR00023 | Alzheimer's disease | 0.013183269 |
| GR00024 | Alzheimer's disease | 0.059322564 |
| GR00025 | Alzheimer's disease | 0.000999421 |
| GR00026 | Alzheimer's disease | 0.013237706 |
| GR00027 | Alzheimer's disease | 0.039542708 |
| GR00028 | Alzheimer's disease | 0.041702778 |
| GR00029 | Alzheimer's disease | 0.03240962 |
| GR00030 | Alzheimer's disease | 0.063687882 |
| GR00031 | Alzheimer's disease | 0.064521106 |
| GR00032 | Alzheimer's disease | 0.047689508 |
| GR00033 | Amyotrophic lateral sclerosis | 0.012234181 |
| GR00034 | Amyotrophic lateral sclerosis | 0.045525479 |

| | | |
|---|---|---|
| GR00035 | Amyotrophic lateral sclerosis | 0.031089518 |
| GR00036 | Anorexia nervosa | 0.019832795 |
| GR00037 | Anti-dsDNA status | 0.067367823 |
| GR00038 | Anti-dsDNA status | 0.204944933 |
| GR00039 | Anxiety disorder | 0.003251616 |
| GR00040 | Aortic valve stenosis | 0.014808642 |
| GR00041 | Arthritis (juvenile idiopathic) | 0.009764383 |
| GR00042 | Arthritis (juvenile idiopathic) | 0.03120038 |
| GR00043 | Asthma | 0.023900759 |
| GR00044 | Asthma | 0.03999514 |
| GR00045 | Asthma | 0.019054731 |
| GR00046 | Asthma | 0.003839564 |
| GR00047 | Asthma | 0.005635514 |
| GR00048 | Asthma | 0.009096351 |
| GR00049 | Asthma | 0.035434514 |
| GR00050 | Atopic dermatitis | 0.008563549 |
| GR00051 | Atopic dermatitis | 0.005507953 |
| GR00052 | Atrial fibrillation | 0.036307888 |
| GR00053 | Atrial fibrillation | 0.127761296 |
| GR00054 | Atrial fibrillation | 0.002327245 |
| GR00055 | Atrial fibrillation | 0.074946298 |
| GR00056 | Autism | 0.006734456 |
| GR00057 | Autism | 0.014552163 |
| GR00058 | Autoimmune hepatitis type-1 | 0.100160246 |
| GR00059 | B cell non-Hodgkin lymphoma | 0.013232706 |
| GR00060 | Barrett's esophagus | 0.004907046 |
| GR00061 | Barrett's esophagus | 0.024962713 |
| GR00062 | Basal cell carcinoma | 0.007403442 |
| GR00063 | Basal cell carcinoma | 0.014777768 |
| GR00064 | Behcet's disease | 0.01485452 |
| GR00065 | Behcet's disease | 0.00551565 |
| GR00066 | Behcet's disease | 0.031025664 |
| GR00067 | Behcet's disease | 0.026699934 |
| GR00068 | Bipolar disorder | 0.045755291 |
| GR00069 | Bipolar disorder | 0.013243625 |
| GR00070 | Black vs. non-black hair color | 0.279026726 |
| GR00071 | Bladder cancer | 0.003302355 |
| GR00072 | Bladder cancer | 0.012165591 |

| | | |
|---|---|---|
| GR00073 | Bladder cancer | 0.002941046 |
| GR00074 | Bladder cancer | 0.034383014 |
| GR00075 | Blond vs non-blond hair color | 0.104755996 |
| GR00076 | Blond vs. brown hair color | 0.07988577 |
| GR00077 | Blue vs. green eyes | 0.012740387 |
| GR00078 | Breast cancer | 0.038264559 |
| GR00079 | Breast cancer | 0.018139919 |
| GR00080 | Breast cancer | 0.006230443 |
| GR00081 | Breast cancer | 0.00456076 |
| GR00082 | Breast cancer | 0.024954645 |
| GR00083 | Breast cancer | 0.008943937 |
| GR00084 | Breast cancer | 0.029914467 |
| GR00085 | Breast cancer | 0.017630407 |
| GR00086 | Breast cancer | 0.009105047 |
| GR00087 | Breast cancer | 0.003570821 |
| GR00088 | Breast cancer | 0.00319403 |
| GR00089 | Breast cancer | 0.003396067 |
| GR00090 | Breast cancer | 0.02029287 |
| GR00091 | Breast cancer | 0.002849713 |
| GR00092 | Breast cancer | 0.048296101 |
| GR00093 | Breast cancer | 0.021239842 |
| GR00094 | Breast cancer | 0.023339868 |
| GR00095 | Breast cancer | 0.056500656 |
| GR00096 | Breast cancer | 0.024113253 |
| GR00097 | Breast cancer | 0.004392492 |
| GR00098 | Breast cancer | 0.05288057 |
| GR00099 | Breast cancer | 0.034816052 |
| GR00100 | Breast cancer | 0.01881074 |
| GR00101 | Breast cancer | 0.00510482 |
| GR00102 | Bronchopulmonary dysplasia | 0.086715224 |
| GR00103 | Brown vs. non-brown hair color | 0.082153762 |
| GR00104 | Brugada syndrome | 0.155369468 |
| GR00105 | Burning and freckling | 0.073277624 |
| GR00106 | Calcific aortic valve stenosis | 0.009144027 |
| GR00107 | Cardia gastric cancer | 0.00524288 |
| GR00108 | Cardiac repolarization | 0.024371607 |
| GR00109 | Cardiovascular disease risk factor | 0.004395567 |
| GR00110 | Cardiovascular disease risk factor | 0.005122735 |

| | | |
|---|---|---|
| GR00111 | Cardiovascular disease risk factor | 0.012372754 |
| GR00112 | Carotid intima media thickness, plaque | 0.002347839 |
| GR00113 | Case-only systemic lupus erythematosus | 0.085437469 |
| GR00114 | Celiac disease | 0.331635197 |
| GR00115 | Cervical cancer | 0.029466815 |
| GR00116 | Cervical cancer | 0.010527487 |
| GR00117 | Cervical cancer | 0.013311215 |
| GR00118 | Cholangiocarcinoma in primary sclerosing cholangitis | 0.016054486 |
| GR00119 | Chronic bronchitis in chronic obstructive pulmonary disease | 0.009217093 |
| GR00120 | Chronic hepatitis B infection | 0.105951642 |
| GR00121 | Chronic hepatitis B infection | 0.118375909 |
| GR00122 | Chronic hepatitis C infection | 0.007928738 |
| GR00123 | Chronic kidney disease | 0.002439075 |
| GR00124 | Chronic kidney disease | 0.00500031 |
| GR00125 | Chronic kidney disease | 0.004448058 |
| GR00126 | Chronic kidney disease | 0.011845313 |
| GR00127 | Chronic kidney disease | 0.014186527 |
| GR00128 | Chronic lymphocytic leukemia | 0.107535678 |
| GR00129 | Chronic lymphocytic leukemia | 0.02346047 |
| GR00130 | Chronic lymphocytic leukemia | 0.034287341 |
| GR00131 | Chronic obstructive pulmonary disease | 0.026073357 |
| GR00132 | Chronic obstructive pulmonary disease | 0.023812372 |
| GR00133 | Chronic obstructive pulmonary disease | 0.027806976 |
| GR00134 | Cleft lip | 0.05404079 |
| GR00135 | Colon cancer | 0.011488937 |
| GR00136 | Colorectal cancer | 0.023427414 |
| GR00137 | Colorectal cancer | 0.010684586 |
| GR00138 | Colorectal cancer | 0.010839925 |
| GR00139 | Colorectal cancer | 0.005617641 |
| GR00140 | Colorectal cancer | 0.020321818 |
| GR00141 | Colorectal cancer | 0.003561416 |
| GR00142 | Colorectal cancer | 0.025715914 |
| GR00143 | Colorectal cancer | 0.020198624 |
| GR00144 | Colorectal cancer | 0.002971312 |
| GR00145 | Colorectal cancer | 0.008986206 |
| GR00146 | Colorectal cancer | 0.045252525 |
| GR00147 | Colorectal cancer | 0.032525407 |

| | | |
|---|---|---|
| GR00148 | Colorectal cancer | 0.017507445 |
| GR00149 | Colorectal cancer | 0.010676326 |
| GR00150 | Colorectal cancer | 0.007081238 |
| GR00151 | Colorectal cancer | 0.010869981 |
| GR00152 | Colorectal cancer | 0.000415944 |
| GR00153 | Colorectal cancer | 0.016858961 |
| GR00154 | Colorectal cancer | 0.008055539 |
| GR00155 | Colorectal cancer | 0.037323325 |
| GR00156 | Combined Crohn's disease and sarcoidosis | 0.004861123 |
| GR00157 | Congenital heart malformation | 0.01166589 |
| GR00158 | Corneal astigmatism | 0.031325538 |
| GR00159 | Coronary artery disease | 0.002338181 |
| GR00160 | Coronary artery disease | 0.012541095 |
| GR00161 | Coronary artery disease | 0.015861535 |
| GR00162 | Coronary artery disease | 0.0324426 |
| GR00163 | Coronary artery disease | 0.011692444 |
| GR00164 | Coronary artery disease | 0.002917435 |
| GR00165 | Coronary artery disease | 0.036175299 |
| GR00166 | Coronary artery disease | 0.005936919 |
| GR00167 | Coronary artery disease | 0.163105336 |
| GR00168 | Coronary artery disease | 0.015613164 |
| GR00169 | Coronary artery disease | 0.02912354 |
| GR00170 | Coronary artery disease | 0.010232528 |
| GR00171 | Coronary artery disease | 0.016804438 |
| GR00172 | Coronary artery disease | 0.004162095 |
| GR00173 | Coronary artery disease | 0.005884303 |
| GR00174 | Coronary artery disease | 0.008804034 |
| GR00175 | Craniofacial microsomia | 0.203447856 |
| GR00176 | Creutzfeldt-Jakob disease | 0.018342418 |
| GR00177 | Crohn's disease | 0.123648489 |
| GR00178 | Crohn's disease | 0.043105554 |
| GR00179 | Crohn's disease | 0.062283135 |
| GR00180 | Crohn's disease | 0.091114789 |
| GR00181 | Crohn's disease | 0.014560295 |
| GR00182 | Crohn's disease | 0.149627553 |
| GR00183 | Curve progression in adolescent idiopathic scoliosis | 0.020970229 |
| GR00184 | Dementia with Lewy bodies | 0.082521523 |
| GR00185 | Diabetic nephropathy | 0.042472652 |

| | | |
|---|---|---|
| GR00186 | Diabetic nephropathy | 0.004488554 |
| GR00187 | Diarrhoeal Disease | 0.00956929 |
| GR00188 | Diarrhoeal Disease | 0.008484939 |
| GR00189 | Diarrhoeal Disease | 0.003612698 |
| GR00190 | Diarrhoeal Disease | 0.006344985 |
| GR00191 | Diffuse large B cell lymphoma | 0.032040324 |
| GR00192 | Digestive system disease | 0.028155603 |
| GR00193 | Disc degeneration (lumbar) | 0.004954629 |
| GR00194 | Disease-free survival in breast cancer | 0.119442865 |
| GR00195 | Drinking behavior | 0.105737067 |
| GR00196 | Drug abuse | 0.015247863 |
| GR00197 | Dupuytren's disease | 0.12598796 |
| GR00198 | Dupuytren's disease | 0.168165597 |
| GR00199 | Early onset inflammatory bowel disease | 0.011425058 |
| GR00200 | Early-onset obesity | 0.041513415 |
| GR00201 | End-stage renal disease in Type 1 diabetics (Women) | 0.02511986 |
| GR00202 | Endometrial cancer | 0.004570587 |
| GR00203 | Endometrial cancer | 0.019239604 |
| GR00204 | Endometrial cancer | 0.019831763 |
| GR00205 | Endometriosis | 0.012651903 |
| GR00206 | Endometriosis | 0.022296848 |
| GR00207 | Endometriosis | 0.017563298 |
| GR00208 | Enteric fever | 0.067629 |
| GR00209 | Epilepsy | 0.021662925 |
| GR00210 | Epilepsy | 0.028071611 |
| GR00211 | Epilepsy | 0.0162159 |
| GR00212 | Epilepsy | 0.019940492 |
| GR00213 | Epithelial ovarian cancer | 0.042557478 |
| GR00214 | Epstein Barr virus nuclear antigen 1 IgG seropositivity | 0.001652333 |
| GR00215 | Esophageal cancer | 0.033814518 |
| GR00216 | Esophageal cancer | 0.029256097 |
| GR00217 | Esophageal cancer | 0.078832239 |
| GR00218 | Esophageal cancer | 0.026815559 |
| GR00219 | Esophageal cancer | 0.016555094 |
| GR00220 | Esophageal cancer | 0.013780689 |
| GR00221 | Esophageal cancer | 0.006138652 |
| GR00222 | Ewing sarcoma | 0.105463706 |

| | | |
|---|---|---|
| GR00223 | Febrile seizures | 0.074429178 |
| GR00224 | Febrile seizures | 0.099680529 |
| GR00225 | Febrile seizures | 0.026919385 |
| GR00226 | Follicular lymphoma | 0.046508738 |
| GR00227 | Follicular lymphoma | 0.028768778 |
| GR00228 | Follicular lymphoma | 0.077641646 |
| GR00229 | Fractures (vertebral) | 0.0148789 |
| GR00230 | Freckles | 0.038379054 |
| GR00231 | Gallbladder cancer | 0.033760555 |
| GR00232 | Gastric cancer | 0.010868997 |
| GR00233 | Glaucoma | 0.026029063 |
| GR00234 | Glaucoma | 0.057532852 |
| GR00235 | Glaucoma | 0.019826004 |
| GR00236 | Glaucoma | 0.018097181 |
| GR00237 | Glaucoma | 0.017521126 |
| GR00238 | Glioma | 0.032164781 |
| GR00239 | Glioma | 0.006910395 |
| GR00240 | Glioma | 0.044742294 |
| GR00241 | Goiter | 0.056663518 |
| GR00242 | Gout | 0.049423696 |
| GR00243 | Gout | 0.137279995 |
| GR00244 | Gout | 0.140782472 |
| GR00245 | Graves disease | 0.137838173 |
| GR00246 | Graves disease | 0.119399435 |
| GR00247 | Graves disease | 0.027304839 |
| GR00248 | Heart failure | 0.166499615 |
| GR00249 | Heart failure | 0.026016022 |
| GR00250 | Helicobacter pylori serologic status | 0.022176897 |
| GR00251 | Hepatitis B | 0.142326493 |
| GR00252 | Hepatitis B | 0.037785111 |
| GR00253 | Hepatitis B | 0.060586149 |
| GR00254 | Hepatitis B | 0.081545902 |
| GR00255 | Hepatitis B | 0.053396341 |
| GR00256 | Hepatitis C induced liver cirrhosis | 0.05108722 |
| GR00257 | Hepatocellular carcinoma | 0.014868406 |
| GR00258 | Hepatocellular carcinoma | 0.029119573 |
| GR00259 | Hepatocellular carcinoma | 0.023644701 |
| GR00260 | Hepatocellular carcinoma | 0.013122643 |

| | | |
|---|---|---|
| GR00261 | High serum lipase activity | 0.035465512 |
| GR00262 | Hodgkin's lymphoma | 0.054789842 |
| GR00263 | Hyperemesis gravidarum | 0.019318486 |
| GR00264 | Hypertension | 0.001824211 |
| GR00265 | Hypertension | 0.008812926 |
| GR00266 | Hypertriglyceridemia | 0.104046998 |
| GR00267 | Idiopathic pulmonary fibrosis | 0.10318996 |
| GR00268 | Idiopathic pulmonary fibrosis | 0.126468724 |
| GR00269 | Immunoglobulin light chain (AL) amyloidosis | 0.075388818 |
| GR00270 | Infantile hypertrophic pyloric stenosis | 0.065542662 |
| GR00271 | Infantile hypertrophic pyloric stenosis | 0.051090279 |
| GR00272 | Inflammatory bowel disease | 0.086864909 |
| GR00273 | Inflammatory bowel disease | 0.103875054 |
| GR00274 | Inguinal hernia | 0.006163864 |
| GR00275 | Insomnia (caffeine-induced) | 0.11860224 |
| GR00276 | Interstitial lung disease | 0.160940486 |
| GR00277 | Intracerebral hemorrhage | 0.010643831 |
| GR00278 | Intracranial aneurysm | 0.030923913 |
| GR00279 | Intracranial aneurysm | 0.019683563 |
| GR00280 | Intracranial aneurysm | 0.031309826 |
| GR00281 | Intracranial aneurysm | 0.020466685 |
| GR00282 | Irritable bowel syndrome | 0.01273299 |
| GR00283 | Ischemic stroke | 0.008444172 |
| GR00284 | Ischemic stroke | 0.003168836 |
| GR00285 | Ischemic stroke | 0.001327427 |
| GR00286 | Ischemic stroke | 0.005851086 |
| GR00287 | Ischemic stroke | 0.003317405 |
| GR00288 | Ischemic stroke | 0.024617484 |
| GR00289 | Kawasaki disease | 0.09915466 |
| GR00290 | Kawasaki disease | 0.016850814 |
| GR00291 | Kidney disease | 0.002010004 |
| GR00292 | Kidney disease | 0.019045123 |
| GR00293 | Kidney disease | 0.011483542 |
| GR00294 | Kidney disease | 0.005471561 |
| GR00295 | Kidney disease | 0.005960596 |
| GR00296 | Kidney stones | 0.004996789 |
| GR00297 | Knee osteoarthritis | 0.012767989 |
| GR00298 | Knee osteoarthritis | 0.010682656 |

| | | |
|---|---|---|
| GR00299 | Late onset Alzheimer's disease | 0.016893913 |
| GR00300 | Late onset Alzheimer's disease | 0.143125219 |
| GR00301 | Leishmaniasis (visceral) | 0.016787029 |
| GR00302 | Leprosy | 0.112480573 |
| GR00303 | Light vs. dark hair color | 0.110772673 |
| GR00304 | Longevity | 0.005771506 |
| GR00305 | Longevity | 0.012461137 |
| GR00306 | Longevity | 0.008818991 |
| GR00307 | Lung adenocarcinoma | 0.020292166 |
| GR00308 | Lung adenocarcinoma | 0.017742513 |
| GR00309 | Lung adenocarcinoma | 0.028713341 |
| GR00310 | Lung adenocarcinoma | 0.016894924 |
| GR00311 | Lung cancer | 0.022623669 |
| GR00312 | Lung cancer | 0.009303033 |
| GR00313 | Lung cancer | 0.000899019 |
| GR00314 | Lung cancer | 0.008773378 |
| GR00315 | Major depressive disorder | 0.021232578 |
| GR00316 | Major depressive disorder | 0.021918913 |
| GR00317 | Major depressive disorder | 0.031924004 |
| GR00318 | Major depressive disorder | 0.008594186 |
| GR00319 | Major depressive disorder | 0.001796614 |
| GR00320 | Major depressive disorder | 0.093204755 |
| GR00321 | Major depressive disorder | 0.113217551 |
| GR00322 | Major depressive disorder | 0.075236357 |
| GR00323 | Major depressive disorder | 0.131593953 |
| GR00324 | Major mood disorders | 0.002518386 |
| GR00325 | Male-pattern baldness | 0.121084438 |
| GR00326 | Marginal zone lymphoma | 0.035517708 |
| GR00327 | Melanoma | 0.001886008 |
| GR00328 | Melanoma | 0.040532491 |
| GR00329 | Meningioma | 0.01859283 |
| GR00330 | Meningioma | 0.006576666 |
| GR00331 | Meningococcal disease | 0.017267286 |
| GR00332 | Mesial temporal lobe epilepsy | 0.005423004 |
| GR00333 | Mesial temporal lobe epilepsy | 0.014282101 |
| GR00334 | Migraine | 0.073711771 |
| GR00335 | Migraine | 0.002755316 |
| GR00336 | Migraine | 0.005754175 |

| | | |
|---|---|---|
| GR00337 | Migraine | 0.033522022 |
| GR00338 | Migraine | 0.0266128 |
| GR00339 | Migraine | 0.016704531 |
| GR00340 | Migraine | 0.042740846 |
| GR00341 | Migraine | 0.050953244 |
| GR00342 | Migraine | 0.035697629 |
| GR00343 | Migraine | 0.016879262 |
| GR00344 | Mitral valve prolapse | 0.042545002 |
| GR00345 | Morning vs. evening chronotype | 0.014771931 |
| GR00346 | Mortality in heart failure | 0.084159468 |
| GR00347 | Mortality in heart failure | 0.039216859 |
| GR00348 | Mortality in heart failure | 0.005810295 |
| GR00349 | Mucinous ovarian carcinoma | 0.037683103 |
| GR00350 | Multiple cancers | 0.005031245 |
| GR00351 | Multiple cancers | 0.002052937 |
| GR00352 | Multiple cancers | 0.004961515 |
| GR00353 | Multiple cancers | 0.004981674 |
| GR00354 | Multiple myeloma | 0.01877838 |
| GR00355 | Multiple myeloma | 0.099548832 |
| GR00356 | Multiple myeloma | 0.047600503 |
| GR00357 | Multiple myeloma | 0.027536656 |
| GR00358 | Multiple myeloma | 0.062994232 |
| GR00359 | Multiple myeloma | 0.125777425 |
| GR00360 | Multiple myeloma | 0.152653675 |
| GR00361 | Multiple myeloma | 0.112659954 |
| GR00362 | Multiple myeloma | 0.040948437 |
| GR00363 | Multiple sclerosis | 0.043567172 |
| GR00364 | Myeloproliferative neoplasms | 0.018935763 |
| GR00365 | Myocardial infarction | 0.031520528 |
| GR00366 | Myopia (pathological) | 0.034021276 |
| GR00367 | Myopic maculopathy | 0.01856891 |
| GR00368 | Narcolepsy | 0.050030281 |
| GR00369 | Narcolepsy | 0.046873759 |
| GR00370 | Nasopharyngeal carcinoma | 0.082867663 |
| GR00371 | Neuroblastoma | 0.01483877 |
| GR00372 | Non-cardia gastric cancer | 0.037716691 |
| GR00373 | Non-cardia gastric cancer | 0.013290882 |
| GR00374 | Non-small cell lung cancer | 0.022057408 |

| | | |
|---|---|---|
| GR00375 | Nonsyndromic cleft lip with or without cleft palate | 0.075608393 |
| GR00376 | Obesity | 0.098733769 |
| GR00377 | Obesity | 0.015721298 |
| GR00378 | Ossification of the posterior longitudinal ligament of the spine | 0.051716579 |
| GR00379 | Osteoarthritis | 0.001672743 |
| GR00380 | Osteoarthritis | 0.000974606 |
| GR00381 | Osteoarthritis | 0.002494576 |
| GR00382 | Osteoarthritis | 0.001845007 |
| GR00383 | Osteoarthritis | 0.004193489 |
| GR00384 | Osteoarthritis | 0.00195496 |
| GR00385 | Osteoarthritis | 0.002922793 |
| GR00386 | Osteoarthritis | 0.018578867 |
| GR00387 | Osteoarthritis | 0.008021126 |
| GR00388 | Osteoarthritis | 0.000645973 |
| GR00389 | Osteoarthritis | 0.000229546 |
| GR00390 | Osteoarthritis | 0.006433369 |
| GR00391 | Osteoarthritis | 0.000127462 |
| GR00392 | Osteoarthritis | 0.010252254 |
| GR00393 | Osteoarthritis | 0.002428762 |
| GR00394 | Osteoporosis | 0.007403442 |
| GR00395 | Osteoporosis | 0.011540439 |
| GR00396 | Ovarian cancer | 0.016626178 |
| GR00397 | Ovarian cancer | 0.005203548 |
| GR00398 | Ovarian cancer | 0.029749261 |
| GR00399 | Ovarian cancer | 0.015756605 |
| GR00400 | Paget's disease of bone | 0.11265716 |
| GR00401 | Pain | 0.003669549 |
| GR00402 | Pancreatic cancer | 0.0088485 |
| GR00403 | Pancreatic cancer | 0.020664923 |
| GR00404 | Pancreatic cancer | 0.004576326 |
| GR00405 | Pancreatic cancer | 0.019372069 |
| GR00406 | Pancreatitis | 0.038531639 |
| GR00407 | Parkinson's disease | 0.023891217 |
| GR00408 | Parkinson's disease | 0.014151241 |
| GR00409 | Parkinson's disease | 0.037396297 |
| GR00410 | Parkinson's disease | 0.02117897 |
| GR00411 | Parkinson's disease | 0.044928409 |

| | | |
|---|---|---|
| GR00412 | Parkinson's disease | 0.065628889 |
| GR00413 | Parkinson's disease | 0.064947411 |
| GR00414 | Parkinson's disease | 0.059609111 |
| GR00415 | Pathological myopia | 0.011414182 |
| GR00416 | Pathological myopia | 0.009392818 |
| GR00417 | Pathological myopia | 0.005383183 |
| GR00418 | Periodontitis | 0.005733699 |
| GR00419 | Peripheral artery disease | 0.005950886 |
| GR00420 | Plantar fascial disorders | 0.000650417 |
| GR00421 | Polycystic ovary syndrome | 0.049405654 |
| GR00422 | Primary biliary cholangitis | 0.108521464 |
| GR00423 | Primary biliary cholangitis | 0.168778978 |
| GR00424 | Primary biliary cholangitis | 0.044073094 |
| GR00425 | Primary biliary cholangitis | 0.063903534 |
| GR00426 | Primary open-angle glaucoma | 0.006113139 |
| GR00427 | Primary sclerosing cholangitis | 0.016860351 |
| GR00428 | Progressive supranuclear palsy | 0.260690268 |
| GR00429 | Progressive supranuclear palsy | 0.00651826 |
| GR00430 | Progressive supranuclear palsy | 0.017385685 |
| GR00431 | Prostate cancer | 0.037336022 |
| GR00432 | Prostate cancer | 0.004880294 |
| GR00433 | Prostate cancer | 0.002755316 |
| GR00434 | Prostate cancer | 0.006591922 |
| GR00435 | Prostate cancer | 0.007694849 |
| GR00436 | Prostate cancer | 0.012092802 |
| GR00437 | Prostate cancer | 0.023114387 |
| GR00438 | Prostate cancer | 0.034007403 |
| GR00439 | Prostate cancer | 0.141671608 |
| GR00440 | Prostate cancer | 0.004087186 |
| GR00441 | Prostate cancer | 0.013263933 |
| GR00442 | Prostate cancer | 0.125502507 |
| GR00443 | Prostate cancer | 0.036620931 |
| GR00444 | Prostate cancer | 0.08792687 |
| GR00445 | Prostate cancer | 0.011899264 |
| GR00446 | Prostate cancer | 0.011526175 |
| GR00447 | Psoriasis | 0.020236227 |
| GR00448 | Psoriasis | 0.033066348 |
| GR00449 | Psoriasis | 0.124705641 |

| | | |
|---|---|---|
| GR00450 | Psoriatic arthritis | 0.075310164 |
| GR00451 | Pulmonary artery enlargement | 0.017126015 |
| GR00452 | Pulmonary artery enlargement | 0.019359247 |
| GR00453 | Red vs non-red hair color | 0.312843634 |
| GR00454 | Red vs. non-red hair color | 0.02780307 |
| GR00455 | Renal cell carcinoma | 0.010859326 |
| GR00456 | Renal cell carcinoma | 0.010229543 |
| GR00457 | Renal gout | 0.111210121 |
| GR00458 | Renal gout | 0.143977121 |
| GR00459 | Response to Dalcetrapib treatment in acute coronary syndrome | 0.027286116 |
| GR00460 | Response to metformin | 0.013311215 |
| GR00461 | Restless legs syndrome | 0.080015375 |
| GR00462 | Restless legs syndrome | 0.050447072 |
| GR00463 | Rhegmatogenous retinal detachment | 0.014978066 |
| GR00464 | Rheumatoid arthritis | 0.005328866 |
| GR00465 | Rheumatoid arthritis | 0.003042565 |
| GR00466 | Rheumatoid arthritis | 0.055045324 |
| GR00467 | Rheumatoid arthritis | 0.055469965 |
| GR00468 | Sarcoidosis | 0.003822456 |
| GR00469 | Sarcoidosis | 0.016984883 |
| GR00470 | Sasang constitutional medicine type (So-Eum) | 0.007287019 |
| GR00471 | Schizophrenia | 0.011696159 |
| GR00472 | Schizophrenia | 0.053531251 |
| GR00473 | Schizophrenia | 0.040260912 |
| GR00474 | Sclerosing cholangitis and ulcerative colitis (combined) | 0.019663288 |
| GR00475 | Scoliosis | 0.007895798 |
| GR00476 | Scoliosis | 0.028621338 |
| GR00477 | Scoliosis | 0.008440551 |
| GR00478 | Shingles | 0.004464138 |
| GR00479 | Sjogren's syndrome | 0.13009326 |
| GR00480 | Skin sensitivity to sun | 0.02780307 |
| GR00481 | Smoking behavior | 0.001048274 |
| GR00482 | Smoking behavior | 0.000342313 |
| GR00483 | Smoking behavior | 0.009165292 |
| GR00484 | Sporadic neuroblastoma | 0.080754199 |
| GR00485 | Sporadic pituitary adenoma | 0.017342659 |
| GR00486 | Squamous cell carcinoma | 0.007006683 |

| | | |
|---|---|---|
| GR00487 | Squamous cell carcinoma | 0.034586255 |
| GR00488 | Stroke | 0.006030095 |
| GR00489 | Stroke | 0.013995513 |
| GR00490 | Stroke | 0.002033565 |
| GR00491 | Stroke | 0.003338235 |
| GR00492 | Stroke | 0.00461742 |
| GR00493 | Stroke | 0.008782398 |
| GR00494 | Stroke | 0.010304228 |
| GR00495 | Stroke | 0.006084844 |
| GR00496 | Sudden cardiac arrest | 0.003558252 |
| GR00497 | Supraventricular ectopy | 0.026160487 |
| GR00498 | Survival in breast cancer | 0.034824375 |
| GR00499 | Survival in colorectal cancer | 0.002909558 |
| GR00500 | Survival in colorectal cancer | 0.004571026 |
| GR00501 | Survival in colorectal cancer | 0.021119802 |
| GR00502 | Susceptibility to persistent hepatitis B virus infection | 0.007381044 |
| GR00503 | Systemic lupus erythematosus | 0.005541145 |
| GR00504 | Systemic lupus erythematosus | 0.009112039 |
| GR00505 | Systemic lupus erythematosus | 0.179247159 |
| GR00506 | Systemic lupus erythematosus | 0.353421906 |
| GR00507 | Systemic lupus erythematosus | 0.017106523 |
| GR00508 | Systemic sclerosis | 0.064608715 |
| GR00509 | Systemic sclerosis | 0.024335964 |
| GR00510 | Systemic sclerosis | 0.124107232 |
| GR00511 | Systemic sclerosis | 0.181583325 |
| GR00512 | Systemic sclerosis | 0.006159744 |
| GR00513 | Systemic sclerosis | 0.012824084 |
| GR00514 | Taxane-induced peripheral neuropathy in breast cancer | 0.181310486 |
| GR00515 | Testicular germ cell cancer | 0.123528788 |
| GR00516 | Testicular germ cell cancer | 0.162714565 |
| GR00517 | Testicular germ cell cancer | 0.024678861 |
| GR00518 | Testicular germ cell cancer | 0.016253323 |
| GR00519 | Thyroid cancer | 0.030497438 |
| GR00520 | Thyroid cancer | 0.078088926 |
| GR00521 | Thyroid cancer | 0.011690141 |
| GR00522 | Tuberculosis | 0.063991801 |
| GR00523 | Tuberculosis | 0.003940065 |

| | | |
|---|---|---|
| GR00524 | Type 1 diabetes | 0.021320715 |
| GR00525 | Type 1 diabetes | 0.051835334 |
| GR00526 | Type 1 diabetes | 0.005037086 |
| GR00527 | Type 1 diabetes nephropathy | 0.00915456 |
| GR00528 | Type 1 diabetes nephropathy | 0.001504595 |
| GR00529 | Type 1 diabetes nephropathy | 0.046037617 |
| GR00530 | Type 2 diabetes | 0.044935806 |
| GR00531 | Type 2 diabetes | 0.001105361 |
| GR00532 | Type 2 diabetes | 0.022416555 |
| GR00533 | Type 2 diabetes | 0.014658754 |
| GR00534 | Type 2 diabetes | 0.054303551 |
| GR00535 | Type 2 diabetes | 0.016754351 |
| GR00536 | Type 2 diabetes | 0.059338112 |
| GR00537 | Type 2 diabetes | 0.027245553 |
| GR00538 | Type 2 diabetes | 0.083002864 |
| GR00539 | Type 2 diabetes | 0.006839448 |
| GR00540 | Type 2 diabetes | 0.004361046 |
| GR00541 | Type 2 diabetes | 0.001610264 |
| GR00542 | Type 2 diabetes | 0.008502074 |
| GR00543 | Type 2 diabetes | 0.0078655 |
| GR00544 | Type 2 diabetes | 0.005614025 |
| GR00545 | Type 2 diabetes | 0.023680631 |
| GR00546 | Type 2 diabetes | 0.023838184 |
| GR00547 | Type 2 diabetes | 0.001134864 |
| GR00548 | Type 2 diabetes | 0.03444121 |
| GR00549 | Type 2 diabetes | 0.006568327 |
| GR00550 | Type 2 diabetes | 0.017650835 |
| GR00551 | Type 2 diabetes | 0.094405465 |
| GR00552 | Type 2 diabetes | 0.041647764 |
| GR00553 | Ulcerative colitis | 0.05540542 |
| GR00554 | Ulcerative colitis | 0.10520555 |
| GR00555 | Ulcerative colitis | 0.061004829 |
| GR00556 | Ulcerative colitis | 0.054964378 |
| GR00557 | Upper aerodigestive tract cancers | 0.01048981 |
| GR00558 | Urinary bladder cancer | 0.009484496 |
| GR00559 | Urinary bladder cancer | 0.013220862 |
| GR00560 | Uterine fibroids | 0.023781525 |
| GR00561 | Uterine fibroids | 0.015926844 |

| | | |
|---|---|---|
| GR00562 | Venous thromboembolism | 0.143738499 |
| GR00563 | Venous thromboembolism | 0.067389334 |
| GR00564 | Venous thromboembolism | 0.082459649 |
| GR00565 | Viral capsid antigen IgG seropositivity | 0.003676288 |
| GR00566 | Vitiligo | 0.009634001 |
| GR00567 | Vitiligo | 0.032379763 |
| GR00568 | Vitiligo | 0.11318408 |
| GR00569 | Wilms tumor | 0.046496324 |
