## Supplementary material for "Assessing the performance of genome-wide association studies for predicting disease risk": S2 Table

| Study Purpose | AUROC | PMID |
|---|---|---|
| Crohn's Disease | 0.75 | 28851283 |
| Alzheimer's Disease | 0.68 | 28727176 |
| Alzheimer's Disease | 0.67 | 26490334 |
| Skin Colour | 0.73-0.97 | 28500464 |
| Pulmonary tuberculosis | 0.64 | 28355295 |
| Breast Cancer | 0.64 | 28205043 |
| Obesity | 0.57 | 23701538 |
| Esophageal Squamous-cell Carcinoma | 0.63 | 23536576 |
| Common Psoriasis | 0.76 | 28537254 |
| Osteoporotic Fracture | 0.55 | 23572424 |
| 10 different conditions | 0.6-0.8 | 28065900 |
| Renal Cell Carcinoma | 0.66 | 27229762 |
| Alcohol Dependence | 0.55 | 23362995 |
| Chronic Myelogenous Leukemia | 0.61 | 26474455 |
| Venous Thromboembolism | 0.66 | 25472531 |
| Alzheimer's Disease | 0.64 | 26543236 |
| Leprosy | 0.71 | 30231057 |
| Colorectal Cancer | 0.59 | 29908285 |
| Head Hair Shape | 0.664-0.789 | 30268682 |
| Celiac Disease | 0.86-0.9 | 24550740 |
| Non-alcoholic Steatohepatitis | 0.65 | 29385134 |
| Type 2 Diabetes | 0.64 | 24947790 |
| Prostate Cancer | 0.60 | 27140652 |
| Crohn's Disease | 0.78 | 27076762 |
| Lung Cancer | 0.55 | 23228068 |
| Triple-Negative Breast Cancer | 0.94 | 28918577 |
| Psoriasis | 0.78 | 28617847 |
| Obesity | 0.73 | 23629956 |
| Trichloroethylene induced hypersensitivity syndrome | 0.81 | 26190474 |
| Bipolar Disorder | 0.60 | 26178159 |
| Obesity in people with major depressive disorder | 0.58 | 25903154 |
| Axial Spondyloarthritis | 0.83 | 27749235 |
| Crohn's Disease | 0.73-0.97 | 28052082 |
| Straight Hair | 0.69 | 26414620 |

| | | |
|---|---|---|
| Nasopharyngeal Carcinoma | 0.52 | 25180181 |
| IVF embryo transfer implantation failures | 0.75 | 28388872 |
| Cutaneous Melanoma | 0.64 | 30060076 |
| Graves' Disease | 0.70 | 30649410 |
| Age Related Macular Degeneration | 0.77 | 24576882 |
| Breast Cancer | 0.63 | 30554720 |
| Osteoporosis | 0.67 | 28580384 |
| Type 2 Diabetes | 0.86 | 29099854 |
| Prostate Biopsy | 0.61 | 24265090 |
| Breast Cancer | 0.57 | 18612136 |
| Type 2 Diabetes | 0.64 | 23956346 |
| Multiple Sclerosis | 0.69 | 22164203 |
| Breast Cancer | 0.59 | 29302764 |
| Non-small Cell Lung Cancer | 0.50 | 23720679 |
| Non-melanoma Skin Cancer | 0.66 | 30085400 |
| Ulcerative Colitis | 0.86 | 24241240 |
| Alzheimer's Disease | 0.70 | 26086184 |
| Rheumatoid Arthritis | 0.79 | 20032229 |
| Rheumatoid Arthritis | 0.66 | 24068971 |
| Prostate Cancer | 0.60 | 26431041 |
| Prostate Biopsy | 0.59 | 22652152 |
| Metabolic Syndrome | 0.64 | 24198294 |
| Breast Cancer | 0.65 | 22269215 |
| Breast Cancer | 0.74 | 23354978 |
| Alzheimer's Disease | 0.69 | 29784544 |
| Effect of Immunosuppressive Treatment in First Kidney Transplant | 0.69 | 27777962 |
| Myocardial Infarcation | 0.60 | 29340220 |
| Rheumatoid Arthritis | 0.59 | 21980439 |
| Non-alcoholic Steatohepatitis | 0.56 | 25597287 |
| Eye color | 0.6-0.889 | 27221533 |
| Breast Cancer | 0.59 | 25380502 |
| Predictors of treatment nonresponse to the first anti-TNF inhibitor in ankylosing spondylitis | 0.77 | 24337767 |
| Hair Color | 0.66-0.86 | 21197618 |
| Breast Cancer | 0.68 | 21212067 |
| Type 2 Diabetes | 0.69 | 30767168 |
| Breast Cancer | 0.58 | 22314178 |
| Alzheimer's Disease | 0.74 | 25720397 |

| | | |
|---|---|---|
| Type 1 Diabetes | 0.92 | 30655379 |
| Systemic Lupus Erythematosus | 0.71 | 29967481 |
| Prostate Cancer | 0.66 | 23071574 |
| Psoriasis | 0.72 | 21559375 |
| Coronary Artery Disease | 0.6-0.8 | 20729558 |
| Breast Cancer | 0.60 | 22585702 |
| Major Depressive Disorder | <0.54 | 25279001 |
| Type 2 Diabetes Mellitus | 0.62 | 20384434 |
| Oral Malignancy | 0.61 | 30657779 |
| Rheumatoid Arthritis | 0.70 | 20309765 |
| Metastic Colorectal Cancer | 0.88 | 25372392 |
| Elevated Liver Fat Content | 0.66 | 23804528 |
| Chronic Hepatitis C Therapy | 0.75 | 23615070 |
| Systemic Lupus Erythematosus | 0.76 | 26689915 |
| Type 1 Diabetes | 0.79 | 19956648 |
| Relapse in recipients of allogeneic haematopoietic stem cell transplantation | 0.72 | 30089915 |
| Breast Cancer | 0.65 | 27279675 |
| Carotid Atherosclerosis | 0.65 | 20941391 |
| Type 2 Diabetes | 0.60 | 18694974 |
| Glaucoma | 0.62-0.94 | 30352225 |
| Type 2 Diabetes | 0.64 | 19197355 |
| Type 2 Diabetes | 0.63 | 20571754 |
| Breast cancer | 0.64 | 27565998 |
| Prostate Cancer | 0.66 | 28827750 |
| Colorectal Cancer | 0.56 | 28233817 |
| Stroke Risk | 0.65 | 30073812 |
| Oral Cancer Recurrence | 0.64 | 28400480 |
| Breast Cancer | 0.58 | 22972951 |
| Multiple Sclerosis | 0.59 | 23903824 |
| Type 1 Diabetes | 0.88 | 26577414 |
| Type 2 Diabetes | 0.58 | 17020404 |
| Type 2 Diabetes | 0.60 | 18591388 |
| Type 2 Diabetes | 0.63 | 19247372 |
| Type 2 Diabetes | 0.60 | 19404609 |
| Type 2 Diabetes | 0.62 | 19862325 |
| Colorectal Cancer | 0.57 | 22490517 |
| Age Related Macular Degeneration | 0.82 | 22666427 |
| Crohn's Disease | 0.71 | 21548950 |

| | | |
|---|---|---|
| Prostate Cancer | 0.61 | 20620408 |
| Age Related Macular Degeneration | 0.73 | 18596911 |
| Age Related Macular Degeneration | 0.77 | 19825847 |
