## Supplementary material for "Assessing the performance of genome-wide association studies for predicting disease risk": S3 Table

| Number of SNPs | Number of Studies in GWAS Central |
|---|---|
| 1 | 165 |
| 2 | 78 |
| 3 | 64 |
| 4 | 54 |
| 5 | 37 |
| 6 | 31 |
| 7 | 19 |
| 8 | 21 |
| 9 | 15 |
| 10 | 20 |
| 11 | 3 |
| 12 | 4 |
| 13 | 4 |
| 14 | 7 |
| 15 | 3 |
| 16 | 4 |
| 17 | 4 |
| 18 | 1 |
| 19 | 1 |
| 20 | 2 |
| 21 | 2 |
| 23 | 4 |
| 25 | 1 |
| 26 | 1 |
| 27 | 1 |
| 28 | 1 |
| 29 | 3 |
| 30 | 3 |
| 32 | 2 |
| 34 | 2 |
| 36 | 2 |
| 37 | 1 |
| 38 | 1 |
| 40 | 1 |
| 44 | 1 |

| | |
|---|---|
| 50 | 6 |
