## Supplementary material for "Assessing the performance of genome-wide association studies for predicting disease risk": S4 Table

| Phenotype/Condition | AUROC |
|---|---|
| Black vs. non-black hair color | 0.97 |
| Celiac disease | 0.84 |
| Dupuytren's disease | 0.75 |
| Age-related macular degeneration | 0.75 |
| Progressive supranuclear palsy | 0.71 |
| Heart failure | 0.66 |
| Testicular germ cell cancer | 0.65 |
| Inflammatory bowel disease | 0.64 |
| Autoimmune hepatitis type-1 | 0.63 |
| Chronic hepatitis B infection | 0.63 |
| Brugada syndrome | 0.63 |
| Blond vs. brown hair color | 0.63 |
| Atrial fibrillation | 0.62 |
| Disease-free survival in breast cancer | 0.62 |
| Taxane-induced peripheral neuropathy in breast cancer | 0.62 |
| Crohn's disease | 0.62 |
| Systemic lupus erythematosus | 0.61 |
| Craniofacial microsomia | 0.61 |
| Paget's disease of bone | 0.61 |
| Leprosy | 0.61 |
| Hypertriglyceridemia | 0.61 |
| Nasopharyngeal carcinoma | 0.61 |
| Anti-dsDNA status | 0.61 |
| Interstitial lung disease | 0.60 |
| Male-pattern baldness | 0.60 |
| Late onset Alzheimer's disease | 0.60 |
| Chronic lymphocytic leukemia | 0.60 |
| Hepatitis B | 0.60 |
| Ulcerative colitis | 0.60 |
| Venous thromboembolism | 0.59 |
| Graves disease | 0.59 |
| Psoriasis | 0.59 |
| Sjogren's syndrome | 0.59 |
| Hodgkin's lymphoma | 0.59 |
| Infantile hypertrophic pyloric stenosis | 0.59 |

| | |
|---|---|
| Primary biliary cholangitis | 0.59 |
| Restless legs syndrome | 0.58 |
| Insomnia (caffeine-induced) | 0.58 |
| Renal gout | 0.58 |
| Burning and freckling | 0.58 |
| High serum lipase activity | 0.58 |
| Prostate cancer | 0.57 |
| Kawasaki disease | 0.57 |
| Idiopathic pulmonary fibrosis | 0.57 |
| Dementia with Lewy bodies | 0.57 |
| Febrile seizures | 0.57 |
| Goiter | 0.57 |
| Sporadic neuroblastoma | 0.57 |
| Gout | 0.57 |
| Polycystic ovary syndrome | 0.57 |
| Early-onset obesity | 0.56 |
| Multiple sclerosis | 0.56 |
| Corneal astigmatism | 0.56 |
| Psoriatic arthritis | 0.56 |
| Immunoglobulin light chain (AL) amyloidosis | 0.56 |
| Nonsyndromic cleft lip with or without cleft palate | 0.56 |
| Blond vs non-blond hair color | 0.56 |
| Supraventricular ectopy | 0.56 |
| Major depressive disorder | 0.56 |
| Ewing sarcoma | 0.56 |
| Glioma | 0.56 |
| Acute-on-chronic liver failure in hepatitis B | 0.56 |
| Bronchopulmonary dysplasia | 0.56 |
| Pancreatitis | 0.56 |
| Myocardial infarction | 0.56 |
| Migraine | 0.56 |
| End-stage renal disease in Type 1 diabetics (Women) | 0.56 |
| Case-only systemic lupus erythematosus | 0.56 |
| Thyroid cancer | 0.56 |
| Hepatitis C induced liver cirrhosis | 0.55 |
| Wilms tumor | 0.55 |
| Gallbladder cancer | 0.55 |
| Light vs. dark hair color | 0.55 |
| Follicular lymphoma | 0.55 |

| | |
|---|---|
| Aggressive periodontitis | 0.55 |
| Acute lymphoblastic leukemia (childhood) | 0.55 |
| Survival in breast cancer | 0.55 |
| Adolescent idiopathic scoliosis | 0.55 |
| Multiple myeloma | 0.55 |
| Myeloproliferative neoplasms | 0.55 |
| Freckles | 0.55 |
| Brown vs. non-brown hair color | 0.55 |
| Lung adenocarcinoma | 0.55 |
| Ossification of the posterior longitudinal ligament of the spine | 0.55 |
| Systemic sclerosis | 0.55 |
| Primary sclerosing cholangitis | 0.54 |
| Parkinson's disease | 0.54 |
| Creutzfeldt-Jakob disease | 0.54 |
| Obesity | 0.54 |
| Response to Dalcetrapib treatment in acute coronary syndrome | 0.54 |
| Leishmaniasis (visceral) | 0.54 |
| Narcolepsy | 0.54 |
| Marginal zone lymphoma | 0.54 |
| Melanoma | 0.54 |
| Rheumatoid arthritis | 0.54 |
| Vitiligo | 0.54 |
| Diffuse large B cell lymphoma | 0.54 |
| Myopic maculopathy | 0.54 |
| Acne (severe) | 0.54 |
| Non-small cell lung cancer | 0.54 |
| Helicobacter pylori serologic status | 0.54 |
| Epithelial ovarian cancer | 0.54 |
| Mucinous ovarian carcinoma | 0.54 |
| Abdominal aortic aneurysm | 0.54 |
| Myopia (pathological) | 0.54 |
| Allergic sensitization | 0.54 |
| Ovarian cancer | 0.54 |
| Red vs. non-red hair color | 0.54 |
| Skin sensitivity to sun | 0.54 |
| B cell non-Hodgkin lymphoma | 0.54 |
| Amyotrophic lateral sclerosis | 0.54 |
| Mortality in heart failure | 0.54 |

| | |
|---|---|
| Cholangiocarcinoma in primary sclerosing cholangitis | 0.54 |
| Meningioma | 0.54 |
| Aortic valve stenosis | 0.53 |
| Cleft lip | 0.53 |
| Allergic disease | 0.53 |
| Endometrial cancer | 0.53 |
| Anorexia nervosa | 0.53 |
| Diabetic nephropathy | 0.53 |
| Response to metformin | 0.53 |
| Neuroblastoma | 0.53 |
| Periodontitis | 0.53 |
| Fractures (vertebral) | 0.53 |
| Mitral valve prolapse | 0.53 |
| Scoliosis | 0.53 |
| Mesial temporal lobe epilepsy | 0.53 |
| Chronic bronchitis in chronic obstructive pulmonary disease | 0.53 |
| Susceptibility to persistent hepatitis B virus infection | 0.53 |
| Digestive system disease | 0.53 |
| Early onset inflammatory bowel disease | 0.53 |
| Chronic hepatitis C infection | 0.53 |
| Autism | 0.53 |
| Gastric cancer | 0.53 |
| Sporadic pituitary adenoma | 0.53 |
| Irritable bowel syndrome | 0.53 |
| Calcific aortic valve stenosis | 0.53 |
| Longevity | 0.52 |
| Coronary artery disease | 0.52 |
| Sclerosing cholangitis and ulcerative colitis (combined) | 0.52 |
| Cardiac repolarization | 0.52 |
| Type 2 diabetes | 0.52 |
| Breast cancer | 0.52 |
| Pulmonary artery enlargement | 0.52 |
| Epilepsy | 0.52 |
| Alzheimer's disease | 0.52 |
| Behcet's disease | 0.52 |
| Knee osteoarthritis | 0.52 |
| Urinary bladder cancer | 0.52 |
| Type 1 diabetes | 0.52 |
| Rhegmatogenous retinal detachment | 0.52 |

| | |
|---|---|
| Non-cardia gastric cancer | 0.52 |
| Bipolar disorder | 0.52 |
| Hepatocellular carcinoma | 0.52 |
| Intracranial aneurysm | 0.52 |
| Squamous cell carcinoma | 0.52 |
| Primary open-angle glaucoma | 0.52 |
| Morning vs. evening chronotype | 0.52 |
| Cardia gastric cancer | 0.52 |
| Congenital heart malformation | 0.52 |
| Pancreatic cancer | 0.52 |
| Inguinal hernia | 0.52 |
| Osteoporosis | 0.52 |
| Hypertension | 0.52 |
| Esophageal cancer | 0.52 |
| Endometriosis | 0.52 |
| Uterine fibroids | 0.52 |
| Disc degeneration (lumbar) | 0.52 |
| Chronic obstructive pulmonary disease | 0.52 |
| Cervical cancer | 0.52 |
| Combined Crohn's disease and sarcoidosis | 0.52 |
| Peripheral artery disease | 0.52 |
| Sarcoidosis | 0.52 |
| Diarrhoeal Disease | 0.52 |
| Anxiety disorder | 0.52 |
| Intracerebral hemorrhage | 0.52 |
| Atopic dermatitis | 0.52 |
| Glaucoma | 0.52 |
| Hyperemesis gravidarum | 0.52 |
| Survival in colorectal cancer | 0.52 |
| Type 1 diabetes nephropathy | 0.52 |
| Pain | 0.51 |
| Basal cell carcinoma | 0.51 |
| Upper aerodigestive tract cancers | 0.51 |
| Tuberculosis | 0.51 |
| Renal cell carcinoma | 0.51 |
| Curve progression in adolescent idiopathic scoliosis | 0.51 |
| Sudden cardiac arrest | 0.51 |
| Drinking behavior | 0.51 |
| Blue vs. green eyes | 0.51 |

| | |
|---|---|
| Major mood disorders | 0.51 |
| Barrett's esophagus | 0.51 |
| Drug abuse | 0.51 |
| Asthma | 0.51 |
| Pathological myopia | 0.51 |
| Cardiovascular disease risk factor | 0.51 |
| Arthritis (juvenile idiopathic) | 0.51 |
| Multiple cancers | 0.51 |
| Smoking behavior | 0.51 |
| Lung cancer | 0.51 |
| Schizophrenia | 0.51 |
| Meningococcal disease | 0.51 |
| Red vs non-red hair color | 0.51 |
| Epstein Barr virus nuclear antigen 1 IgG seropositivity | 0.50 |
| Viral capsid antigen IgG seropositivity | 0.50 |
| Sasang constitutional medicine type (So-Eum) | 0.50 |
| Kidney disease | 0.50 |
| Kidney stones | 0.50 |
| Plantar fascial disorders | 0.50 |
| Shingles | 0.50 |
| Colon cancer | 0.50 |
| Carotid intima media thickness, plaque | 0.50 |
| Enteric fever | 0.50 |
| Colorectal cancer | 0.50 |
| Chronic kidney disease | 0.50 |
| Bladder cancer | 0.50 |
| Ischemic stroke | 0.50 |
| Stroke | 0.49 |
| Osteoarthritis | 0.47 |
